# The effects of temporal continuities of grasslands on the diversity and species composition of plants

**DOI:** 10.1101/2020.04.21.050013

**Authors:** Inoue Taiki, Yaida A. Yuki, Uehara Yuki, Katsuhara R. Koki, Kawai Jun, Takashima Keiko, Ushimaru Atushi, Kenta Tanaka

**Affiliations:** Sugadaira Research Station, Mountain Science Center, University of Tsukuba, 1278-294 Sugadaira kogen, Ueda city, Nagano 386-2204, Japan; Graduate School of Human Development and Environment, Kobe University, 3-11 Tsurukabuto, Kobe 657-8501, Japan

**Keywords:** land-use history, ski run, historical land use, semi-natural grassland, endangered plant species

## Abstract

Semi-natural grasslands are ecosystems rich in biodiversity. However, their decline has been reported worldwide, and identification of grasslands with high conservation priority is urgently required. Recently, an increasing number of studies have reported that past vegetation history affects current biological communities. To evaluate whether the temporal continuity of grasslands promotes biodiversity, and thus can be an indicator of conservation priority, we studied vascular plant communities in old (160– 1000s years) and new (52–70 years after deforestation) grasslands, as well as in forests, of Sugadaira Highland in central Japan. The number of plant species was highest in old grasslands, followed by new grasslands and forests. This pattern was much clearer in the number of grassland-dependent native and grassland-dependent endangered species, indicating the role of old grasslands as refugia for those species. The species composition differed between old and new grasslands. New grasslands had species compositions in between those of old grasslands and forests, suggesting that the plant community in new grasslands retains the influence of past forestation for more than 52 years after deforestation. Eleven indicator species were detected in old grasslands, but none in new grasslands, suggesting the uniqueness of the plant community in old grasslands. We conclude that the temporal continuity of grasslands increases plant diversity and can be an indicator of grasslands with high conservation priority.

## Introduction

Grasslands may be natural, existing under natural climatic conditions and disturbance regimes, or they may be semi-natural, whereby they are maintained by artificial disturbances (Squires, Dengler, Feng, & Hua, 2018). Semi-natural temperate grasslands have high species diversity and conservation values (Akeroyd & Page, 2011; Organisation for Economic Co-operation and Development [OECD], 2008; Wilson, Peet, Dengler, & Pärtel, 2012). However, approximately half of the natural grasslands in the world have been converted to farmlands (Goldewijk, 2001), and semi-natural grasslands have also experienced a worldwide decline. In parts of Britain, 47% of semi-natural grasslands disappeared between 1960 and 2013 (Ridding, Redhead, & Pywell, 2015). In Sweden, 38.3% of pasture declined from 1980 to 2003 (Food and Agriculture Organization of the United Nations [FAO], 2010). In Japan, although grasslands constituted 13% of land area 100 years ago, this area has gradually decreased and in the early 2000s grasslands made up just 1% of land area (Ogura, 2006). The decrease in grasslands in Japan is mainly attributed to the decline in the economic value of semi-natural grasslands and the abandonment of their management since 1950, followed by natural succession to forests (Nishiwaki, 1999; Ushimaru, Uchida, & Suka, 2018). This decline in semi-natural grassland area is one of the major causes of biological extinction in Japan (Ministry of the Environment of Japan, 2012, 2016).

Given the rapid decline in semi-natural grasslands due to ongoing social and economic issues, it’s not practical to conserve all of the remaining grasslands. If it was possible to predict which grasslands harbour more biodiversity-rich communities compared to others, high conservation priority could be given to certain ‘hotspot grasslands’. One factor that might be relevant to the formation and retention of biodiversity-rich communities is the history of the vegetation community. The effect of history on species richness and community assembly is one of the central issues in recent research in the field of community ecology. For example, present-day plant species diversity of European grasslands has been shown to be affected by habitat connectivity 50–100 years ago (Lampinen, Heikkinen, Manninen, Ryttäri, & Kuussaari, 2018; Lindborg & Eriksson, 2004). In addition, current butterfly species richness in tropical rainforests has been shown to be affected by past disturbance (Whitworth, Villacampa, Brown, Huarcaya, Downie, & MacLeod, 2016), and bird community composition in a forest is affected by the maintenance of that forests temporal continuity (Culbert, Dorresteijn, Loos, Clayton, Fischer, & Kuemmerle, 2017). Finally, the diversity of arbuscular mycorrhizal fungi has been shown to be higher in grasslands maintained over 12–20 years compared to those maintained for only ten years (Honnay, Helsen, & Van Geel, 2017). Thus, investigating the effect of vegetation history on current grassland plant species biodiversity will not only improve understanding of the process by which biological communities are formed, but also provide valuable knowledge for allocating conservation priority.

Semi-natural grasslands in the Sugadaira Highland of central Japan are believed to have been in a continuous state of existence since 4000 years ago, estimated by ^14^C dating of the underlying Andosol (Yamanoi, 1996), which is known to be specifically generated by and accumulated in grasslands (Yamane, 1973). Since the 1930s, parts of these grasslands have been used as ski runs, and during the last 100 years large parts of these have changed to forests due to forest plantation as well as natural succession triggered by the abandonment of human grassland management. When skiing became popular in Japan in the 1960s–1970s, some parts of these forest plantations were then clear-cut and converted to ski runs. However, overall approximately 92% of grassland area in this region has been converted to forests in the past 100 years. Recently, Yaida, Nagai, Oguro, Katsuhara, Uchida, and Kenta (2019) reported that ski runs in this area functioned as refugia for grassland-specific endangered plants. In this study, we investigated plant communities in old grasslands (160–1000s years) and new grasslands (52–70 years after deforestation), both of which are managed as ski runs in the Sugadaira Highland. Although many studies have only focused on the effect of temporal continuity on species richness (Cousins & Eriksson, 2001, 2002; Gustavsson, Lennartsson, & Emanuelsson, 2007; Johansson et al., 2008), this study dealt with both species richness and composition. In addition, this study includes nearby forests. In order to examine the effect of temporal continuity of grasslands on plant communities and its relevance to conservation priority, this study addressed whether (i) plant diversity was higher in old grasslands compared to new ones; (ii) species composition differed between old and new grasslands; and (iii) the plant communities of new grasslands are still influenced by past forestation.

## Methods

### Study area and sites

Field surveys were conducted in ski run grasslands and adjacent secondary forests in the Sugadaira and Minenohara Highlands of central Japan (latitude 36.51182– 36.55727°, longitude 138.30424–138.35716°, altitude 1319–1522 m) in 2017. Within this area, grasslands were identified and geographical information system (GIS) data generated, using topographic maps from 1910 and 1937, and aerial photographs from 1947, 1975 and 2010. All of these resources are available from the Geospatial Information Authority of Japan (GIAJ, http://mapps.gsi.go.jp/maplibSearch.do#1). An old map (Sugadaira kaikon no zu) from 1855 was also used. In 1855 almost the entire study area, was semi-natural grasslands, probably maintained by pasturing, fire and mowing. Some grasslands have been consistently maintained since that time and have been used as ski runs since 1923. These grasslands, with a long history of the same vegetation (at least 162 years), were defined as old grasslands for this study. Other grasslands were afforested by plantation after 1910, then subsequently deforested between 1947 and 1965 to create ski runs. These grasslands were defined as new grasslands (52 to 70 years old). All ski run grasslands (new and old) are maintained by mowing in the autumn, a conventional form of management (Yaida et al., 2019).

### Field investigation

Within the study area, seven old grasslands, six new grasslands, and seven forests were chosen so that study sites of each vegetation type were distributed as uniformly as possible (Figure 1, Appendix 1). The forest sites were set just beside either old or new grassland sites. All grassland sites were in ski areas, except for one old grassland that was set in the University of Tsukuba’s Sugadaira Research Station as a reference semi-natural grassland free from the effect of skiing. A 1 × 20 m transect was positioned in each study site at least 20 m away from borders between grasslands and forests. In each transect we recorded the presence of vascular plant species in July and September 2017. Several plants were classified into grassland-dependent native species according to Satake, oi, Kitamura, Watari, and Tominari (1981, 1982a, 1982b), and grassland-dependent endangered species by defining endangered species as those above the Near Threatened (NT) rank in the Red lists of at least one prefecture in Japan.

**Figure 1.**
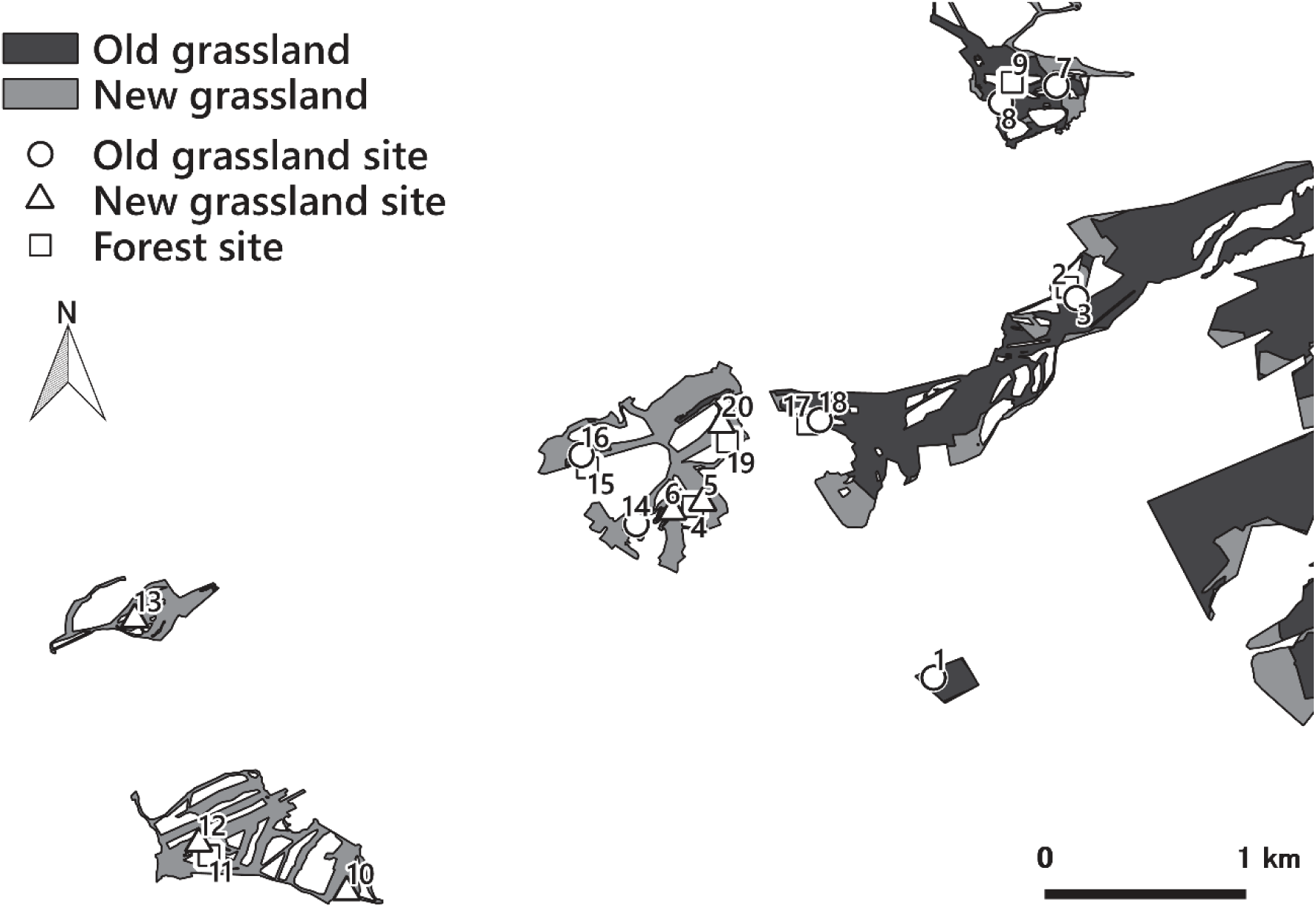
Distribution of old and new grasslands in the Sugadaira and Minenohara Highlands of central Japan. The vegetation type at each study site is indicated by the three symbols (circle = old grassland, triangle = new grassland, and cross = forest). The numbers indicate study sites corresponding to Appendix 1.

### Statistical analyses

The numbers of total plant species, grassland-dependent native species, and grassland-dependent endangered species were compared among vegetation types, using generalized linear models (GLMs) with Poisson distributions and the log link, in the statistics software *R* ver. 3.5.0 (R Core Team, 2018). The following three models were constructed: the null model — the numbers of species are the same among all vegetation types; the two vegetation model — grasslands and forests have different species numbers; and the three vegetation model — old grasslands, new grasslands and forests have different species numbers. Models were selected based on Akaike’s information criterion (AIC) followed by the likelihood ratio test. To graphically ordinate the variation in plant species composition among the vegetation types, non-metric multidimensional scaling (NMDS) with the Jaccard index as the index of dissimilarity was conducted using the *vegan* library (Oksanen et al., 2018) in *R*. A permutational multivariate analysis of variance (PERMANOVA) (Anderson, 2001; McArdle & Anderson, 2001) was used to test the effect of grassland type (old or new) on the species composition. This analysis was implemented using the *adonis* function in the *vegan* library (Oksanen et al., 2018) in *R*, with 10,000 permutations. A Mantel test was conducted, using the *vegan* library in *R*, to detect whether there was spatial autocorrelation between the dissimilarity index and geographic distance for all pairs of grassland sites. Indicator species were detected for each of the three vegetation types using the *indval* function (IndVal test; see Dufrêne & Legendre, 1997) in the *labdsv* library (Roberts, 2016) in *R*. The IndVal test was repeated for the three-vegetation-type case (old grasslands, new grasslands and forests), two-vegetation-type case (grasslands and forests), and one-vegetation-type case (vegetation type not discriminated), as recommended by Dufrêne & Legendre (1997). For each species the indicator value was compared across each vegetation type for each case. Each species was then regarded as an indicator for the vegetation type in which it had the highest indicator value.

## Results

In total, 245 plant species were detected. For total plant species richness the three vegetation model was selected, indicating that each vegetation type effects plant diversity differently (ΔAIC to null model = 53.5, GLM and likelihood ratio test: *p* < 0.001, Table 1a). The number of plant species in each transect was the highest in old grasslands (n=7), followed by new grasslands (n=6) and forests (n=7) (Figure 2). The difference between old and new grasslands was supported for all vascular plants (ΔAIC to second model = 3.8, GLM and likelihood ratio test: *p* < 0.05, Figure 2, Table 1a). Differences in the species richness of grassland-dependent native plants among vegetation types (ΔAIC to the second model = 20.1, GLM and likelihood ratio test: *p* < 0.001, Figure 3, Table 1b) and that in grassland-dependent endangered plants (ΔAIC to the second model = 14.3, GLM and likelihood ratio test: *p* < 0.001, Figure 4, Table 1c) were larger than that in species richness of all plant species.

**Table 1.** Model selection for GLMs regarding the effects of vegetation types on species richness of all (a), grassland-dependent native (b), and grassland-dependent endangered (c), vascular plants. Forests, old grasslands and new grasslands were discriminated for the three vegetation model, and forests and grasslands for the two vegetation model. All vegetation types were treated as the same for the null model. Bold indicates the best model. Delta AIC shows the difference in AIC between the best model and the others. *p* values are for the likelihood ratio test between the best model and each of the others. d.f. indicates degrees of freedom.

| Model | d.f. | AIC | ΔAIC | $p$ |
| --- | --- | --- | --- | --- |
| (a) Species richness of all vascular plants |  |  |  |  |
| **Three vegetation model** | 3 | 238.3 | 0.0 |  |
| Two vegetation model | 2 | 242.3 | 4.0 | < 0.05 |
| Null model | 1 | 292.3 | 54.0 | < 0.001 |
| (b) Species richness of grassland-dependent native vascular plants |  |  |  |  |
| **Three vegetation model** | 3 | 168.1 | 0.0 |  |
| Two vegetation model | 2 | 188.2 | 20.1 | < 0.001 |
| Null model | 1 | 326.7 | 158.6 | < 0.001 |
| (c) Species richness of grassland-dependent endangered vascular plants |  |  |  |  |
| **Three vegetation model** | 3 | 150.0 | 0.0 |  |
| Two vegetation model | 2 | 164.3 | 14.3 | < 0.001 |
| Null model | 1 | 250.1 | 100.1 | < 0.001 |

**Figure 2.**
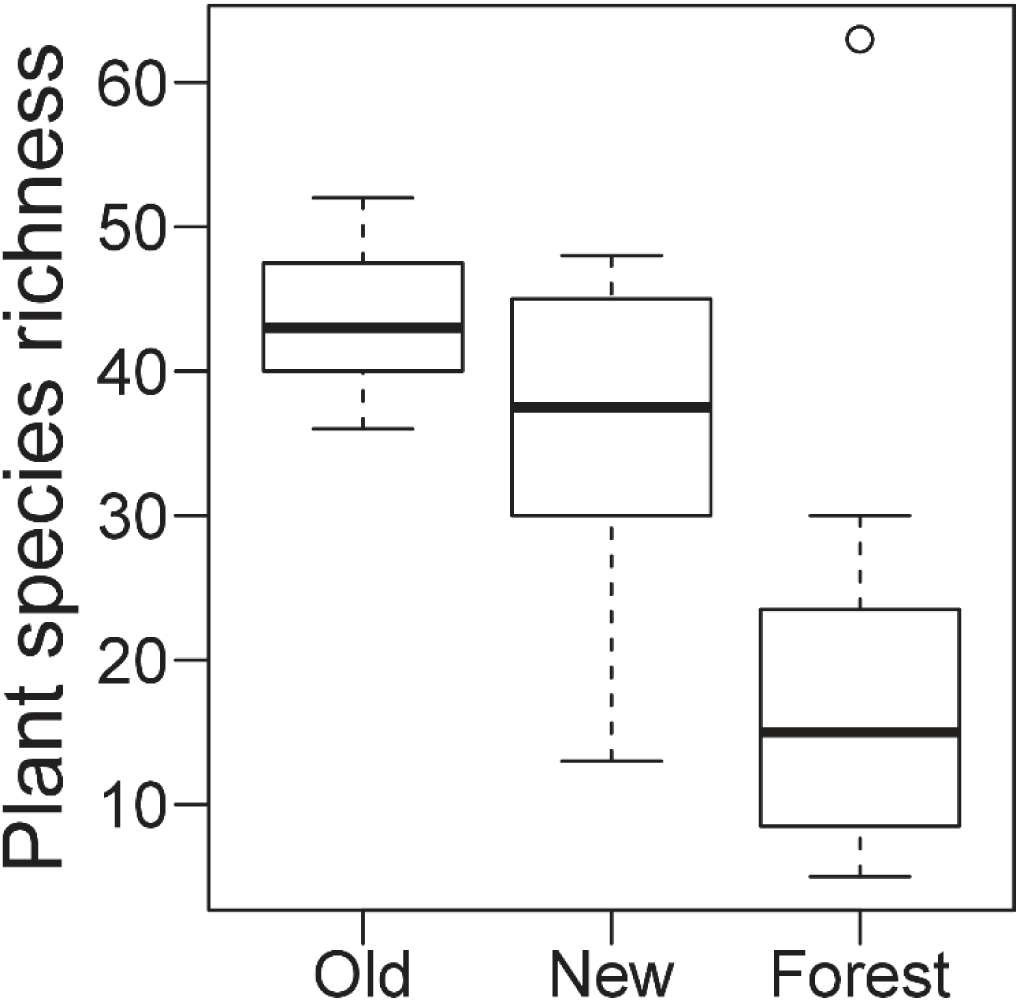
The number of plant species in each vegetation type. “Old” and “New” refer to old and new grasslands (see detail in the text). Bars and boxes are medians and quartiles, with ranges between minimum and maximum values. An outlier (values less than the first quartile minus 1.5 times the interquartile range (IQR) or greater than the third quartile plus 1.5 times the IQR) is shown by a dot.

**Figure 3.**
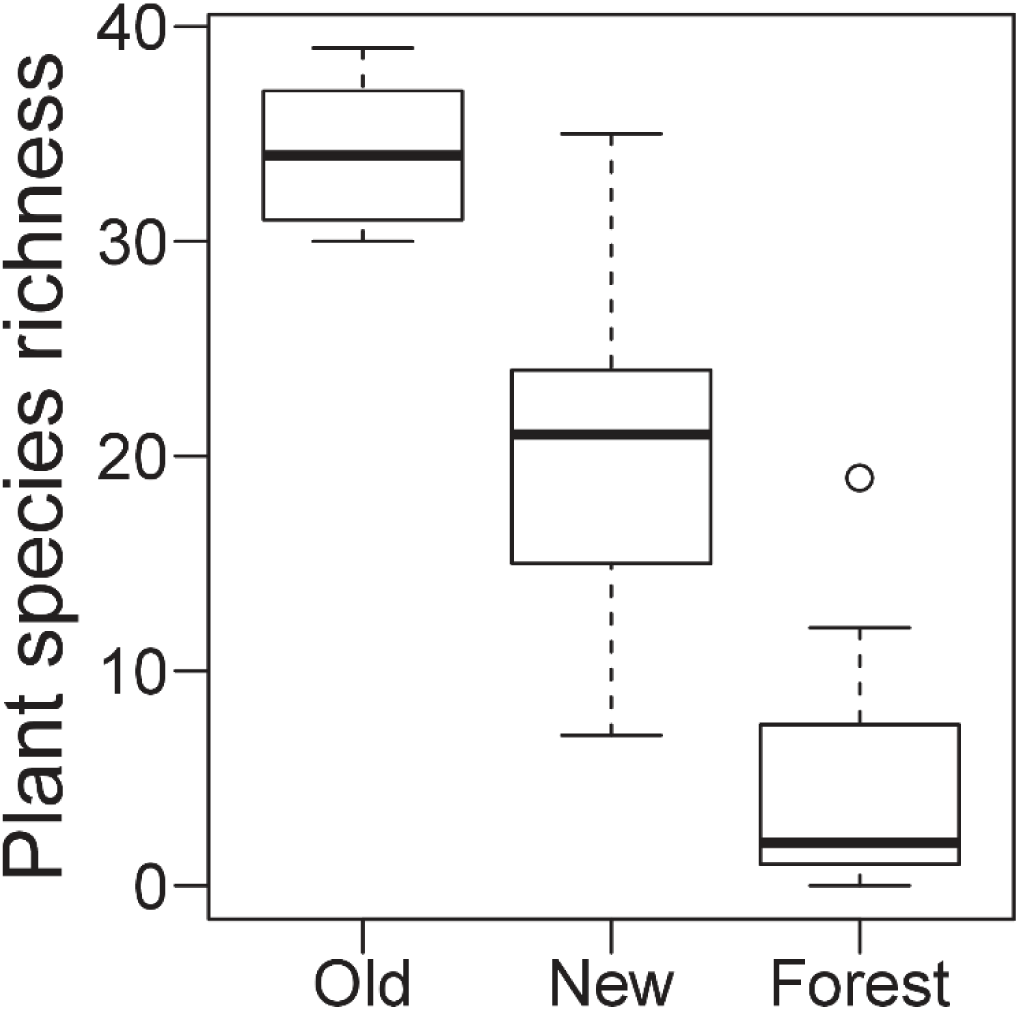
The number of grassland-dependent native plant species found in each vegetation type. Bars and boxes are medians and quartiles, with ranges between minimum and maximum values. Outliers are shown by dots.

**Figure 4.**
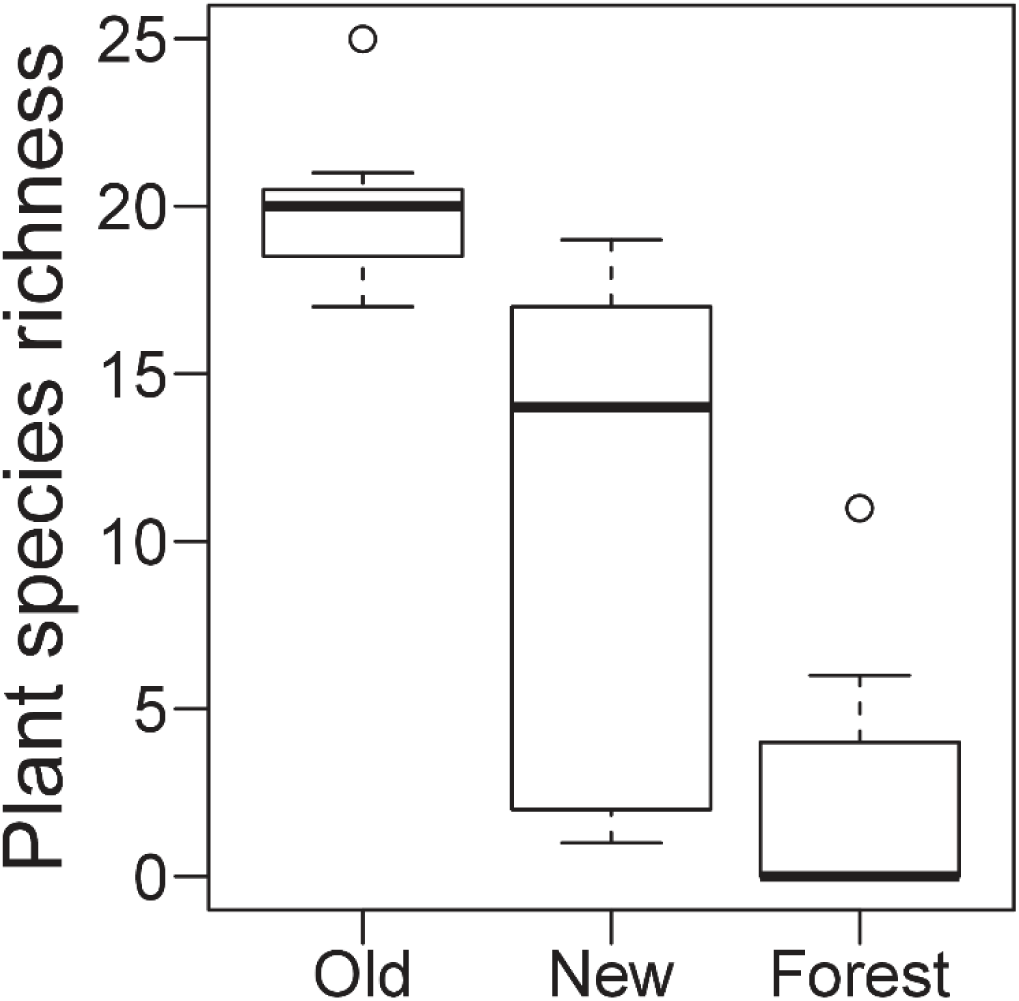
The number of grassland-dependent endangered plant species found in each vegetation type. Bars and boxes are medians and quartiles, with ranges between minimum and maximum values. Outliers are shown by dots.

Plant species compositions significantly differed among vegetation types, both between grasslands and forests, and between old and new grasslands (PERMANOVA: *p* < 0.001 and *p* < 0.01). Old grassland sites were narrowly distributed within the NMDS plot (Figure 5), showing that old grasslands have a specific vegetation community. New grassland sites were distributed between old grassland and forest sites, showing that the species composition is more similar between new grasslands and forests than between old grasslands and forests. The relationship between dissimilarity calculated by the Jaccard index and spatial distance is shown in Figure 6. Spatial autocorrelation in plant species composition was not detected (Mantel test, *p* = 0.139).

**Figure 5.**
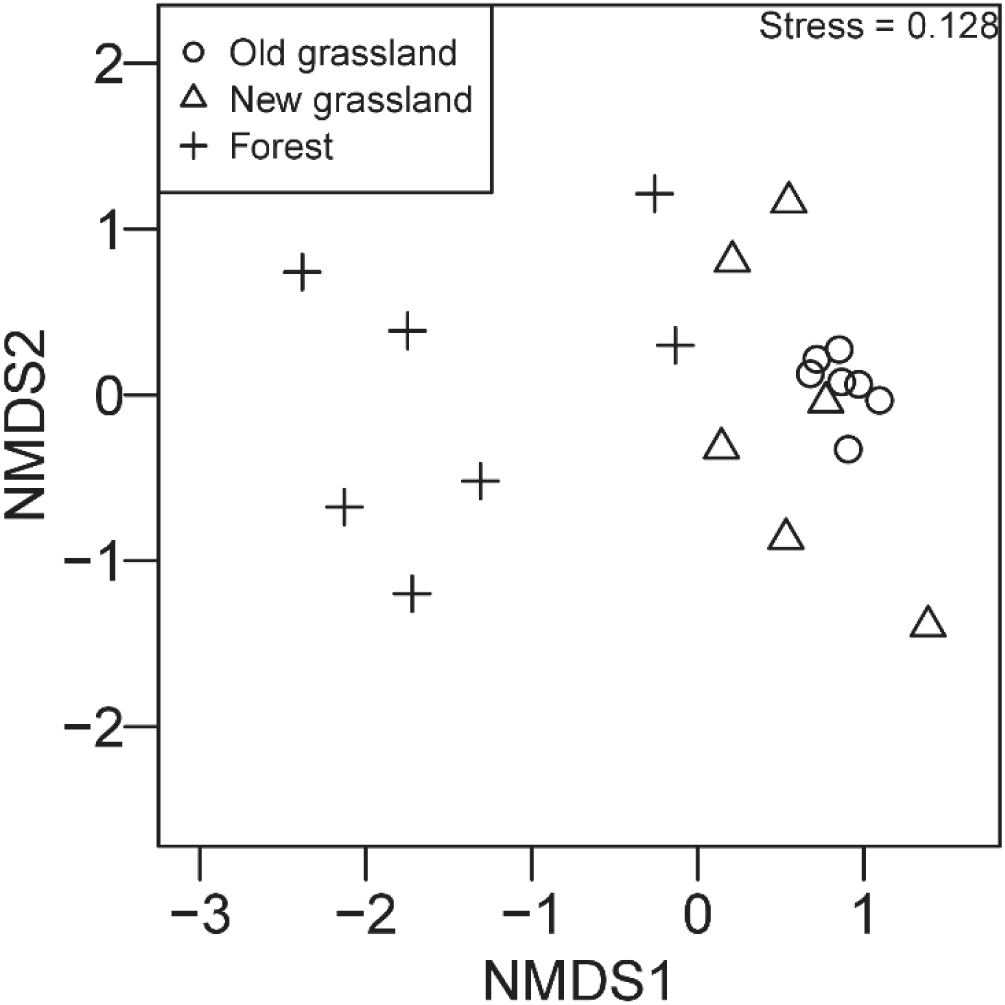
NMDS (non-metric multidimensional scaling) plot using the Jaccard index of dissimilarity of the plant composition of each study site across the three vegetation types.

**Figure 6.**
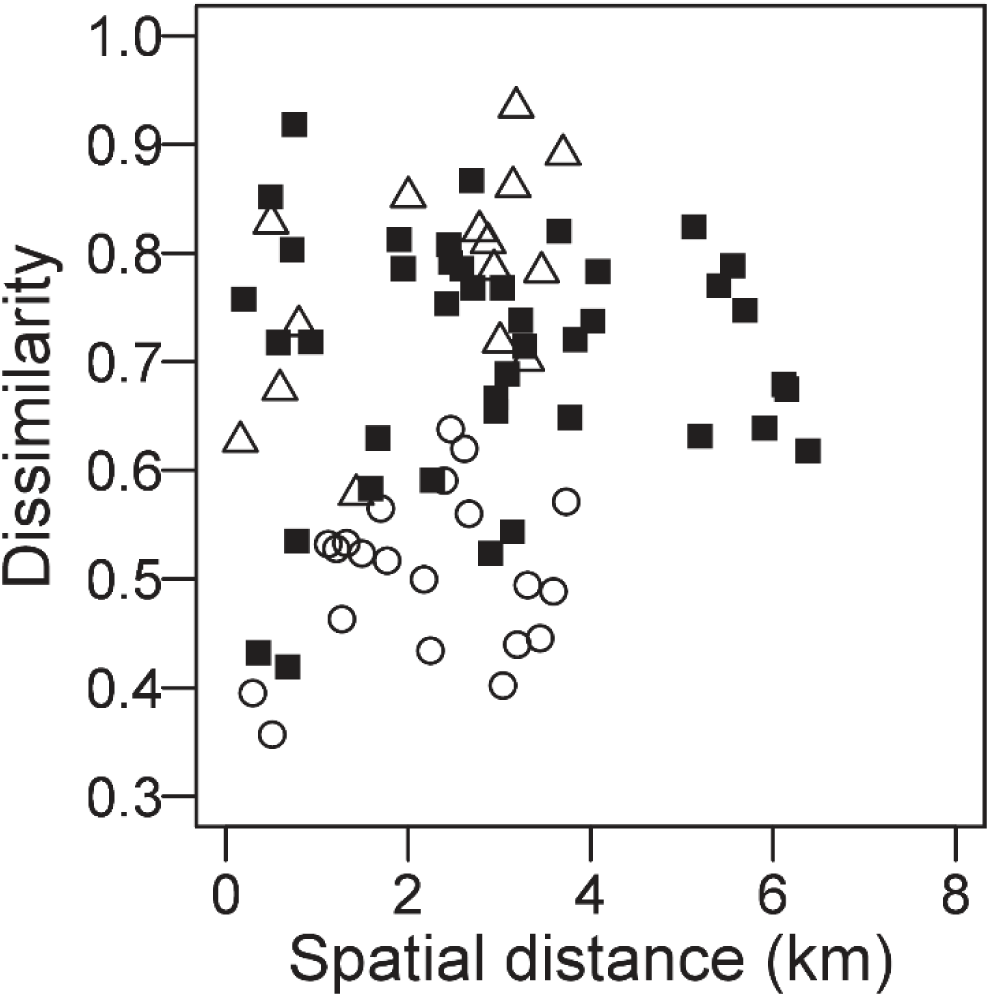
The relationship between the dissimilarity calculated by the Jaccard index and geographical distance for all pairs of grassland sites: old grassland vs old grassland (open circle), new grassland vs new grassland (open triangle), and old grassland vs new grassland (filled square).

The IndVal test detected 11 indicator plant species for old grasslands, seven for grasslands (including both old and new), and two for forests (Table 2). However, no species was considered an indicator of new grasslands (Table 2).

**Table 2.** Results of the indicator species analysis. Only species with significant indicator values are shown (*p* < 0.05), except for those in the indicator group ‘All’.

| Species | Indicator group | Indicator value | $p$-value |
| --- | --- | --- | --- |
| *Rubus subcrataegifolius* | All | 65 | 1.000 |
| *Celastrus orbiculatus* | All | 65 | 1.000 |
| *Solidago virgaurea* | All | 55 | 1.000 |
| *Convallaria majalis* | All | 50 | 1.000 |
| *Viola grypoceras* | All | 50 | 1.000 |
| *Artemisia indica* | Grassland | 92 | 0.001 |
| *Miscanthus sinensis* | Grassland | 72 | 0.006 |
| *Potentilla freyniana* | Grassland | 72 | 0.003 |
| *Iris sanguinea* | Grassland | 69 | 0.007 |
| *Angelica pubescens* | Grassland | 69 | 0.008 |
| *Pteridium aquilinum* | Grassland | 65 | 0.018 |
| *Arenaria lateriflora* | Grassland | 65 | 0.022 |
| *Toxicodendron orientale* | Forest | 57 | 0.005 |
| *Maackia amurensis* | Forest | 43 | 0.032 |
| *Toxicodendron trichocarpum* | Forest | 43 | 0.042 |
| *Adenophora triphylla* | Old grassland | 100 | 0.001 |
| *Sanguisorba officinalis* | Old grassland | 100 | 0.001 |
| *Polygonatum odoratum* | Old grassland | 74 | 0.002 |
| *Cirsium* spp. | Old grassland | 72 | 0.001 |
| *Carex nervata* | Old grassland | 72 | 0.001 |
| *Lespedeza bicolor* | Old grassland | 68 | 0.003 |
| *Galium verum* | Old grassland | 62 | 0.007 |
| *Prunella vulgaris* | Old grassland | 58 | 0.016 |
| *Hosta sieboldiana* | Old grassland | 57 | 0.019 |
| *Lysimachia clethroides* | Old grassland | 55 | 0.021 |
| *Ixeridium dentatum* | Old grassland | 50 | 0.040 |

## Discussion

Compared with new grasslands, higher plant diversity was found in old grasslands (Figure 2), where the temporal continuity of grasslands was longer. These findings are in accordance with a previous study that reported a higher number of plant species in older grasslands (Gustavsson et al., 2007) compared to newly generated grasslands that were once used as arable lands. The current study also showed that old grasslands have a role as refugia for endangered species that depend on grasslands (Figure 4). The finding that the diversity of grassland-dependent native species and grassland-dependent endangered species were clearly higher in the old grasslands (Figure 3, 4), is in agreeance with previous studies (Gustavsson et al., 2007; Johansson et al., 2008; Waldén, Öckinger, Winsa, & Lindborg, 2017) reporting richer grassland specialist species in old grasslands. Because the positive effect of the temporal continuity of a grassland on plant diversity has only been previously reported in Sweden (Gustavsson et al., 2007), the current study (in Japan) suggests the generality of this effect independent of other factors such as continent. In addition, the current findings suggest that plant diversity can be negatively affected by discontinuity of grassland history due to both cultivation (Gustavsson et al., 2007) and forestation (this study). However, some previous studies (Cousins & Eriksson, 2002; Culbert et al., 2017; Johansson et al., 2008) did not detect a difference in plant diversity between old and new grasslands. These studies might contain several sources of heterogeneity; such as regional heterogeneity due to a large study area (approximately 90 km) in Culbert et al. (2017), soil heterogeneity including wet, mesic, and dry soil in Cousins and Eriksson (2002), and management heterogeneity stemming from different levels of discontinuity in Johansson et al. (2008). Higher levels of heterogeneity increase the variance of species richness values even within the same grassland type; so that the effect of the temporal continuity of a grassland may be more difficult to detect. In contrast, this study, and that of Gustavsson et al. (2007), examined the effect within a narrow spatial-scale, using a simple binary comparison between continuity and discontinuity, and for only mesic or dry soil conditions.

This study confirmed that plant species composition differs between old and new grasslands (Figure 5). This finding concurs with a study by Honnay et al. (2017), which examined grasslands of different ages (8–20 years old). However, the current study covered a much longer time-scale, revealing that 52–70 years old grasslands had different plant communities from the older ones (160–1000s years old). In contrast to grassland sites, forest sites had a larger variation in species composition, shown in the NMDS plot (Figure 5). The Jaccard similarity index, which corresponds to the number of shared species between sites, was low between all forest sites. Also, a fewer number of indicator species were detected in forest sites (Table 2). Combined, these results suggest that plant communities vary considerably between forest sites, which can probably, at least partly, be explained by the fact that several different types of forests were included in this study (*Larix* plantation and *Quercus*-dominated forests with several types of forest floor). Despite such variation, the NMDS plot still showed that all of the forest sites formed a single cluster separated from both types of grasslands. However, the plant species composition of new grasslands and forests was more similar than that of old grasslands and forests (Figure 5), suggesting that new grasslands still harbor plant species that recruited in forests. Previous studies (Jonason, Bergman, Westerberg, & Milberg, 2016; Jonason et al., 2014) have reported that plant communities in grasslands generated after clearing forests are influenced by whether those forests once experienced grassland vegetation before forestation. Jonason et al. (2014) showed that grassland species remained as remnant populations after the afforestation. However, this study detected few grassland plant species in the forest sites (Figure 3,4), implying that even a single event of forestation eliminates many grassland plants. Although all of the new grasslands used to be semi-natural grasslands with a long history, many plant species, especially grassland-dependent ones, did not restore viable populations. Such irreversible effects of forestation found in this study have important implications for conserving grassland biodiversity.

Eleven indicator species were detected for the old grasslands, but none for the new grasslands (Table 2), suggesting that the old grasslands have a cluster of unique species as shown in Figure 5, whereas the new grasslands do not have particular characteristics in their species composition. This pattern can be explained by seed-dispersal limitation during the colonization of newly-generated grasslands (Bischoff, 2002; Rosenthal, 2006). Whilst plants with low seed-dispersal ability are most likely limited to old grasslands, those with high seed-dispersal ability could be distributed in both the old and new grasslands. This pattern was most clearly pronounced in *Adenophora triphylla* and *Sanguisorba officinalis* (indicator values of 100, Table 2): both of these species were present in all seven old grasslands but were absent in all the other sites (seven new grasslands and six forests). The high dependence of these two species on old grasslands can also be explained by their low seed-dispersal ability as they lack any organs specialized for seed dispersal (Numata, Asano, & Kuwabara, 1990; Numata & Asano, 1969). A study in a lowland plain in eastern Japan (Koyanagi, Kusumoto, Yamamoto, Ohkuro, Ide, & Takeuchi, 2007) also detected these two species as indicator species for grasslands with high plant diversity, implying that those species-rich lowland grasslands may have long temporal continuity that has enabled the two species to recruit.

This study showed that old grasslands had higher plant species diversity, higher richness of grassland-dependent endangered plant species, and unique species compositions, clearly indicating that they are of high conservation priority. Although new grasslands are likely to be in a state of constant formation and disappearance, old grasslands are consistently decreasing because they are irreversibly lost due to the abandonment of grassland followed by forestation, leading endangered species to further decline. Ski runs in Japan are regularly maintained by mowing and include old grasslands with high conservation priority, thus this study further confirmed the important role of ski runs in Japan as refugia for grassland-specific endangered species (Yaida et al., 2019). Given the rapid and large-spatial-scale decline of semi-natural grasslands in Japan as well as across the world, it is not practical to conserve all grasslands. Thus, it is proposed that grasslands with long temporal continuity have high conservation priority and are where conservation effort should be concentrated.

## vii. Acknowledgments

We are grateful to the land owners of our study sites: “general foundation Nirei kai”, “Sugadaira Bokujo”, “Hatsunekan”, “Imai-kan” and “Jozan-kan”, as well as the management companies: “Sugadaira Pine Beak Ski”, “Sugadaira Ski House Co., LTD”, “Oku Davos Snow Park”, and “HARE Sugadaira-Kogen Snow Resort”, for allowing our field surveys. This work was supported by JSPS KAKENHI Grant Number 17K07557.

