## Appendix 1 for "The effects of temporal continuities of grasslands on the diversity and species composition of plants"

### xi. Appendices

Appendix 1. Location of the study sites and environmental factors

| No. | Site name | Type | Longitude (°E) | Latitude (°N) | Slope angle (°) | Elevation (m) | Vegetation height (cm) |
|---|---|---|---|---|---|---|---|
| 1 | Sugadaira research station | Old | 138.34906 | 36.52383 | 5 | 1329 | 92.7 |
| 2 | Oku Davos forest | forest | 138.35657 | 36.54572 | 10 | 1360 | 89.2 |
| 3 | Oku Davos grassland | old | 138.35706 | 36.54511 | 8 | 1378 | 66.8 |
| 4 | Tengu forest | forest | 138.33557 | 36.53343 | 20 | 1359 | 100.3 |
| 5 | Tengu grassland | new | 138.33613 | 36.53364 | 20 | 1352 | 68.6 |
| 6 | Hinode grassland | new | 138.33444 | 36.53315 | 23 | 1344 | 73.4 |
| 7 | Minenohara B grassland | old | 138.35595 | 36.55701 | 17 | 1472 | 96.4 |
| 8 | Minenohara C grassland | old | 138.35284 | 36.55607 | 17 | 1412 | 78.8 |
| 9 | Minenohara forest | forest | 138.35347 | 36.55721 | 20 | 1433 | 24.7 |
| 10 | Omatsu east grassland | new | 138.31618 | 36.51183 | 22 | 1463 | 87.3 |
| 11 | Omatsu forest | forest | 138.30846 | 36.51387 | 26 | 1522 | 132.5 |
| 12 | Omatsu grassland | new | 138.30788 | 36.51454 | 28 | 1514 | 112.1 |
| 13 | Tsubakuro grassland | new | 138.30429 | 36.52710 | 31 | 1426 | 117.5 |
| 14 | Omotetaro grassland | old | 138.33241 | 36.53244 | 26 | 1319 | 104.3 |
| 15 | Shirogane forest | forest | 138.32967 | 36.53560 | 12 | 1327 | 88.6 |
| 16 | Shirogane grassland | old | 138.32933 | 36.53631 | 14 | 1324 | 107.5 |
| 17 | Ura Davos forest | forest | 138.34201 | 36.53805 | 24 | 1360 | 111.5 |
| 18 | Ura Davos grassland | old | 138.34267 | 36.53827 | 9 | 1378 | 37.1 |
| 19 | Urataro forest | forest | 138.33751 | 36.53706 | 16 | 1354 | 33.1 |
| 20 | Urataro grassland | new | 138.33717 | 36.53803 | 16 | 1339 | 99.0 |
